# Maximizing PHB content in *Synechocystis sp.* PCC 6803: development of a new photosynthetic overproduction strain

**DOI:** 10.1101/2020.10.22.350660

**Authors:** Moritz Koch, Jonas Bruckmoser, Jörg Scholl, Waldemar Hauf, Bernhard Rieger, Karl Forchhammer

## Abstract

PHB (poly-hydroxy-butyrate) represents a promising bioplastic variety with good biodegradation properties. Furthermore, PHB can be produced completely carbon-neutral when synthesized in the natural producer cyanobacterium *Synechocystis sp.* PCC 6803. This model strain has a long history of various attempts to further boost its low amounts of produced intracellular PHB of ~15 % per cell-dry-weight (CDW).

We have created a new strain that lacks the regulatory protein PirC (gene product of *sll0944*), which causes a rapid conversion of the intracellular glycogen pools to PHB under nutrient limiting conditions. To further improve the intracellular PHB content, two genes from the PHB metabolism, phaA and phaB from the known production strain *Cupriavidus necator*, were introduced under the regime of the strong promotor P*psbA2*. The created strain, termed PPT1 (Δ*sll0944*-RE*phaAB*), produced high amounts of PHB under continuous light as well under day-night rhythm. When grown in nitrogen and phosphor depleted medium, the cells produced up to 63 % / CDW. Upon the addition of acetate, the content was further increased to 81 % / CDW. The produced polymer consists of pure PHB, which is highly isotactic.

The achieved amounts were the highest ever reported in any known cyanobacterium and demonstrate the potential of cyanobacteria for a sustainable, industrial production of PHB.

## 1. Introduction

The global contamination with non-degradable plastic is a huge environmental burden of our time (Jambeck et al., 2015, Li et al., 2016). While bioplastics have been suggested as a potential solution, they still represents only a very small fraction of the overall used plastics (Geyer et al., 2017). Furthermore, many of these bioplastics have unsatisfying biodegradation properties. The most common bioplastic, PLA (poly-lactic-acid), is almost undegradable in marine environments (Narancic et al., 2018). This led to the emerging interest in another class of bioplastics with improved degradation properties: PHAs (poly-hydroxy-alkanoates). The most common variant in this chemical class is PHB (poly-hydroxy-butyrate) which is produced by various microorganisms. Currently, PHB is produced by fermentation using heterotrophic bacteria, such as *Cupriavidus necator* or *Escherichia coli (Chen, 2009)*. However, since these production processes require crop-derived organic carbon sources for growth and production, it conflicts with human food-supply. An alternative way of producing PHB, which is independent of cropland use, is the usage of phototrophic organisms, such as cyanobacteria (Balaji et al., 2013, Akiyama et al., 2011). *Synechocystis sp.* PCC 6803 (hereafter *Synechocystis*) is a well-studied model organism for phototrophic growth and a natural producer of PHB (Hein et al., 1998, Wu et al., 2001). Under conditions of nutrient limitation, for example nitrogen starvation, the cells transform into a resting state during a process that is called chlorosis (Allen and Smith, 1969). During chlorosis, they do not only degrade their photosynthetic apparatus, but also accumulate large quantities of glycogen as a carbon- and energy-storage (Klotz et al., 2016, Doello et al., 2018). During the later stages of chlorosis, the cells start to degrade glycogen and convert it to PHB (Koch et al., 2019). However, the produced amounts of intracellular PHB are rather low and only range between 10 - 20 % / CDW (cell dry weight). A recent economic analysis suggests that one of the limiting factors to compete with PHB derived from fermentative processes is the low ratio of PHB / CDW in cyanobacteria (Knöttner et al., 2019). One major goal is therefore, to optimize cyanobacteria so that they achieve higher intracellular PHB contents. This would not only increase the yield but would also simplify the downstream-process of extracting the PHB from the cells.

There have been various attempts to further boost the amount of PHB in cyanobacterial cells. A selection of important approaches has been listed in **Table 1**.

**Table 1.** Previous attempts to optimized the medium or genetic background of *Synechocystis sp*. PCC 6803 with for the production of PHB. Further approaches (also in other cyanobacteria) have been reviewed recently (Kamravamanesh et al., 2018b).

| Genotype | PHB content (% CW) | Substrate | Production condition | Polymer composition | Reference |
| --- | --- | --- | --- | --- | --- |
| WT | 29 | 0.4 % acetate | -P | PHB | (Panda et al., 2006) |
| overexpression *phaAB* (native) | 35 | 0.4 % acetate | -N | PHB | (Khetkorn et al., 2016a) |
| overexpression *phaABC* (*Cupriavidus necator*) | 11 | 10 mM acetate | -N | PHB | (Sudesh et al., 2002) |
| overexpression *nphT7*, *phaB*, *phaC* | 41 | 0.4 % acetate | Limited air exchange, -N | - | (Lau et al., 2014) |
| overexpression *Xfpk* | 12 | CO<sub>2</sub> | -N, -P | PHB | (Carpine et al., 2017) |
| overexpression *sigE* | 1.4 | CO<sub>2</sub> | -N | PHB | (Osanai et al., 2013) |
| overexpression *rre37* | 1.2 | CO<sub>2</sub> | -N | PHB | (Osanai et al., 2014) |

Most of the attempts in the past have focused on genetical engineering approaches to reroute the intracellular flux towards PHB (Carpine et al., 2017, Lau et al., 2014, Osanai et al., 2013, Osanai et al., 2014). *Synechocystis* is naturally producing PHB from acetyl-CoA via the enzymes acetyl-CoA acetyltransferase (PhaA), acetoacetyl-CoA reductase (PhaB) and the heterodimeric PHB synthase (PhaEC). The overproduction of the genes encoding for those enzymes is known for increasing the PHB content within the cells (Khetkorn et al., 2016a, Sudesh et al., 2002).

The highest reported rate of photosynthetically produced PHB in a wildtype (WT) cyanobacterium was achieved in a strain isolated from a wet volcanic rock in Japan. This strain, *Synechococcus sp.* MA19, achieved 27 % / CDW (Miyake et al., 1996). It has to be mentioned though that other groups, who tried to work with this strain, reported that they were unable to obtain it from any known strain collection or laboratory (Markl et al., 2018). Hence, it has to be assumed that this strain disappeared. Another mentionable approach was achieved by applying UV radiation for random mutagenesis (Kamravamanesh et al., 2018a). The created *Synechocystis sp*. PCC 6714 strain produced up to 37 % PHB / CDW under phototrophic growth with CO_2_ as the sole carbon source.

Besides genetical engineering, applying optimized growth conditions and medium was demonstrated to also increase PHB production (Panda et al., 2006). A study investigating 137 different cyanobacterial species found that 88 of them produced PHB, depending on the nutrient, which was lacking in the growth medium (Kaewbai-Ngam et al., 2016). The highest yields were often achieved when cells were starved for nitrogen. Furthermore, the addition of organic carbon sources, like acetate or fructose, resulted in increased PHB production (Panda et al., 2006). A comprehensive overview about different approaches can be found in recent reviews (Kamravamanesh et al., 2018b, Singh and Mallick, 2017). Conflicting results concerning PHB synthesis were reported from attempts, where cells were grown under conditions of limited gas exchange. Whereas some groups reported increased yields (Panda et al., 2006, Lau et al., 2014), other groups reported that they were unable to reproduce this effect (Kamravamanesh et al., 2017). Furthermore, a recent study demonstrated that cells, which were grown under standing-conditions and were thereby also exposed to limited gas-exchange, exhibited a decreased PHB accumulation (Koch et al., 2020). Despite these various approaches to further increase the PHB content in *Synechocystis,* the highest PHB contents reached so far are still far beyond what was accomplished in heterotrophic bacteria, where more than 80 % of biomass is converted into the desired product.

We have recently identified a gene, sll0944, whose deletion resulted in significantly increased PHB synthesis. The Sll0944 (“PirC”) deficient mutant converted its intracellular glycogen pool under nitrogen starvation rapidly to PHB (Orthwein et al., 2020). Therefore, the aim of this study was to create a strain with maximized PHB content by combining the sll0944 mutation with other factors that improve PHB synthesis. This resulted in a strain that can accumulate more than 80% PHB, which is by far the most efficient PHB producing oxygenic photosynthetic organism reported to date.

## 2. Results

In this study, we wanted to test if the PHB content of a mutant strain based on a Δ*sll0944* background can be further increased. Recently it was shown, that overexpression of the PHB synthase PhaEC in *Synechocystis* PCC 6803 can cause a reduction of the PHB production, while overexpression of its *phaAB* genes caused an increase in intracellular PHB accumulation (Khetkorn et al., 2016a). Here, we cloned and overexpressed *phaA* and *phaB* from the known PHB production strain *Cupriavidus necator* (formerly known as *Ralstonia eutropha*) into a Δ*sll0944* strain. We used these genes, since they are derived from an highly efficient PHB synthesizing organism. Furthermore, the expression of heterologous enzymes ensures that these enzymes are not inhibited by intracellular regulatory mechanisms. Both genes were cloned into a pVZ322 vector under the regime of a strong promotor P*psbA2*. The plasmid was then transformed into the strain Δ*sll0944*, thereby creating the strain Δ*sll0944*-RE*phaAB* (Figure S1). For simplifications, the strain was termed PPT1 (for PHB Producer Tübingen 1).

### Strain characterization

To compare the growth of the newly generated strain with the WT, both strains were grown under continuous illumination as well as under a 12/12 hours light/dark regime (**Figure 1**).

**Figure 1.**
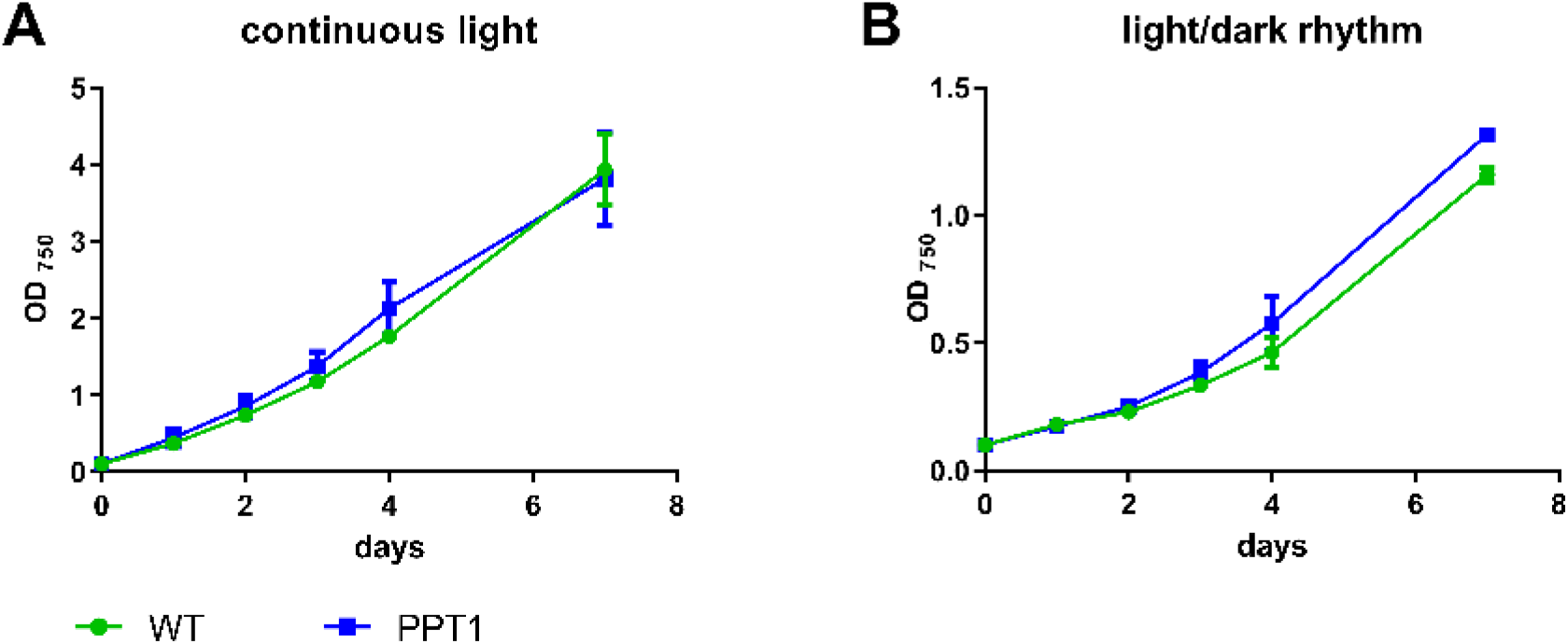
Growth behavior of WT and PPT1 strains grown under continuous illumination (A) or under a 12/12 hour light/dark regime (B). The growth was determined over 7 days by recording the OD_750_. Each point represents a mean of three independent biological replicates.

Under both light regimes, WT and PPT1 strain exhibited similar growth rates. This was also the case when the strains were grown on solid agar plates (Figure S2). To test whether the mutant strain was able to produce PHB under vegetative growth, the PHB production of both strains was tested in BG_11_ medium during exponential and stationary growth stages (OD_750_ ~1 and ~3, respectively) (Figure S3). While the WT did not produce any detectable amounts of PHB under exponential growth, the mutant accumulated ~ 0.5 % / CDW. Under stationary conditions, none of both strains produced any detectable amount of PHB.

To test, whether the newly generated mutant is able to accumulate higher amounts of PHB under production conditions, different cultivation conditions were systematically tested. The conditions of the highest production rates were then used for further experiments. First, the impact of continuous illumination compared to day-night cycles was tested. Therefore, WT and PPT1 cells were shifted to nitrogen-free BG_0_ medium to induce chlorosis and were subsequently grown under 12/12 hours light /dark cycle or under continuous illumination; the amount of intracellular PHB was quantified and normalized to the CDW (Figure 2). For an easier comparison, all following graphs about PHB accumulation have the same y-axis scalation.

**Figure 2.**
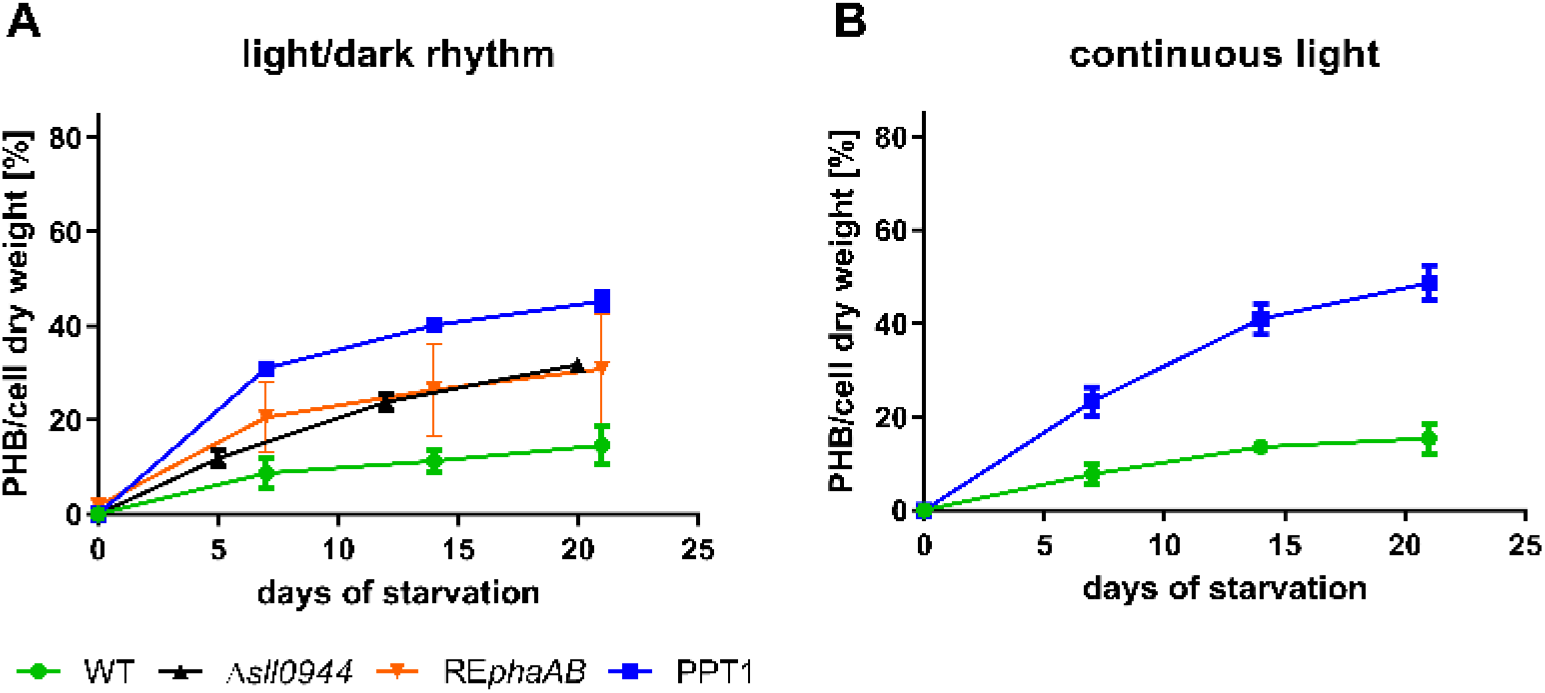
PHB content of WT (blue), Δ*sll0944* (black), RE*phaAB* (orange) and PPT1 (green) cells with different light regimes. Exponentially grown cells were shifted to nitrogen free BG_0_ and cultivated under either diurnal (12 hours light/12 hours darkness) (A) or continuous light (B). Each point represents a mean of three independent biological replicates.

To test the influence of the individual genetic modifications, the PHB content of two strains harbouring only one of the two genetic alterations (Δ*sll0944* or RE*phaAB*, respectively) were measured. Compared to the WT, the Δ*sll0944* and the RE*phaAB* strains produced higher amounts of PHB (32 and 31 % / CDW, respectively) after three weeks of chlorosis. When both mutations were combined (PPT1), the accumulation of PHB was further increased to 48 and 45 % PHB / CDW at dark/light or continuous light, respectively. With 31 % of PHB per CDW after 7 days in diurnal cultivation, the initial rate of PHB synthesis in the PPT1 cells was higher as compared to continuous illumination, where PHB amounted to 23 %. Therefore, these conditions were further investigated.

### Medium optimization

Other studies have reported that, besides nitrogen, the lack of other elements can also induce the biosynthesis of PHB in *Synechocystis* (Kaewbai-Ngam et al., 2016). To test this effect on the newly generated strain, WT and PPT1 cells were shifted to either sulphur, phosphor or nitrogen/phosphor-free medium and the content of intracellular PHB was quantified (Figure 3).

**Figure 3.**
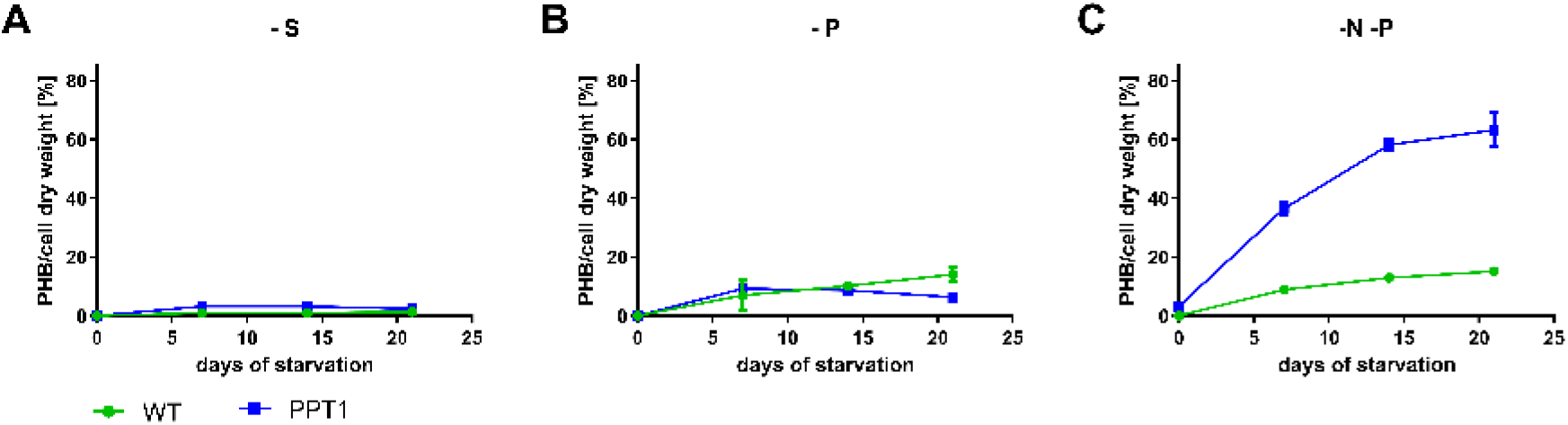
PHB content of WT (green) and PPT1 (blue) cells grown in different media under dark/light rhythm. To induce PHB production, exponentially grown cells were shifted to either sulphur, phosphor or nitrogen/phosphor free medium (A, B and C, respectively). Each point represents a mean of three independent biological replicates.

Whenever phosphate free production conditions were used, the precultures were already grown in phosphate-free BG_11_, in order to deplete the intracellular polyphosphate storage pools of *Synechocystis*. In sulphur-as well as in phosphor-free medium, both strains produced only minor amounts of PHB. However, when the cells were shifted to nitrogen/phosphor-free medium, the mutant strain accumulated after three weeks high amounts of up to 63 % / CDW. Under the same conditions, the WT accumulated only 15 % / CDW. All further experiments will be based on cultures grown in nitrogen- and phosphor depleted BG_11_ medium.

To test, if the produced PHB amounts can be further increased by the addition of an additional carbon sources, either 100 mM NaHCO_3_ or 10 mM acetate were added after the shift to nitrogen/phosphor-free medium (Figure 4).

**Figure 4.**
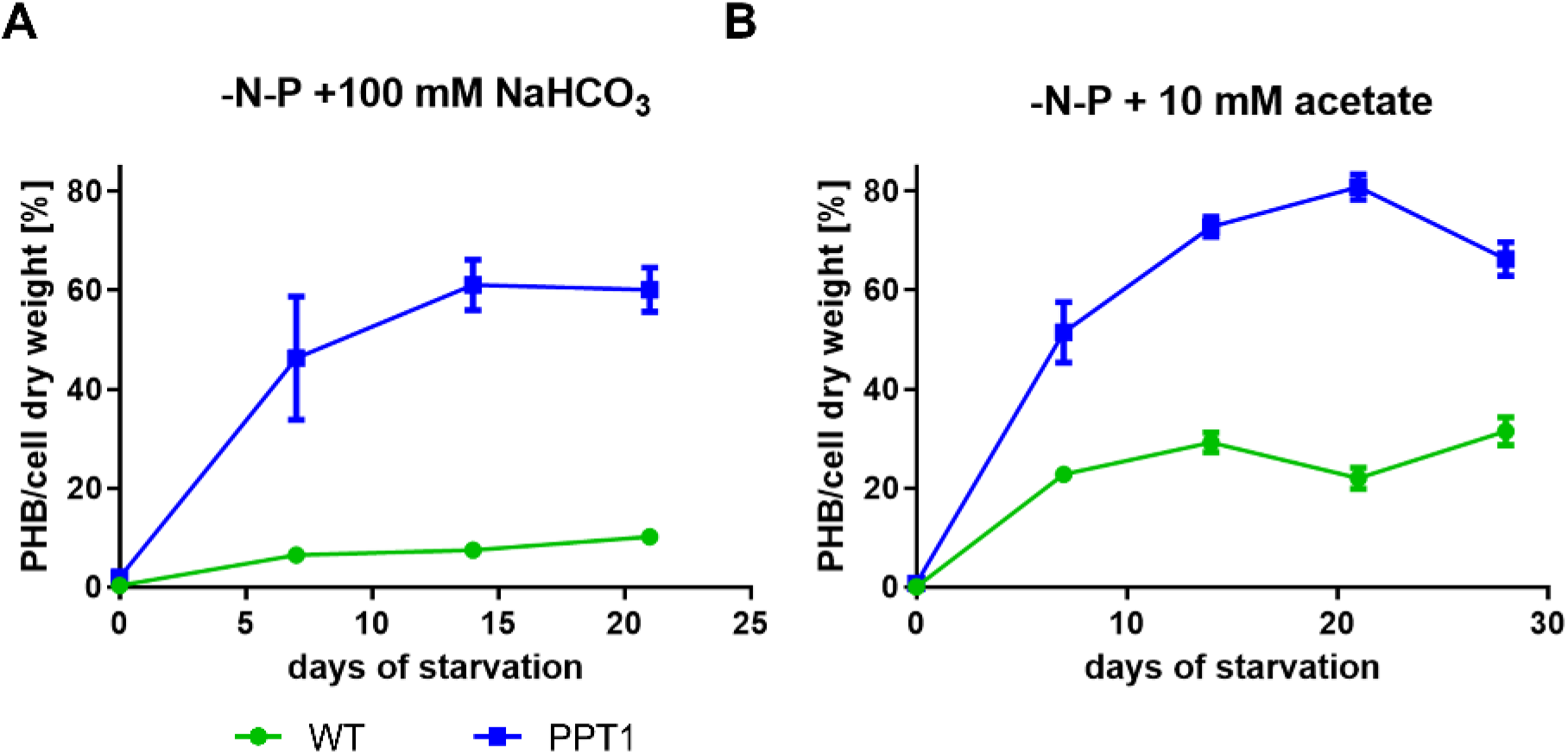
PHB production of WT (green) and PPT1 (blue) cells grown under alternating light/dark regime. (A) Cells shifted to nitrogen/phosphor free medium with the addition of 100 mM NaHCO_3_. (B) Cells shifted to nitrogen/phosphor free medium with the addition of 10 mM acetate. Each point represents a mean of three independent biological replicates.

As in the previous experiments, cells were again cultivated in diurnal light/dark regime. When NaHCO_3_ was added, the PPT1 cells reached intracellular PHB contents of up to 61 % / CDW after two weeks, while the WT accumulated only 10 % / CDW. Notably, the initial production rate was further increased, leading to an average of 46 % / CDW in the PPT1 after one week. When instead of NaHCO_3_ 10 mM acetate were added, the WT reached intracellular PHB contents of up to 32 % / CDW after four weeks, while the PPT1 mutant accumulated up to 81 % / CDW after three weeks of starvation (Figure 4 A). A further starvation of another week did not further increase the yields, but instead slightly reduced the intracellular amount of PHB. When cells were grown under the same conditions but with continuous illumination, the produced amounts of PHB were much lower (Figure S4).

To test if the limitation of gas-exchange could lead to a further increase of PHB production, nitrogen-phosphorus starved cells were grown in sealed vessels. However, after an initial increase of intracellular PHB, the amount dropped strongly (Figure S5).

### Visualization of PHB granules

To find out how the high PHB values that were quantified by HPLC analysis affect the morphology of the cells, and how these masses of PHB are organized within the cells, fluorescence microscopy as well as transmission-electron-microscopy (TEM) pictures were taken (Figure 5).

**Figure 5.**
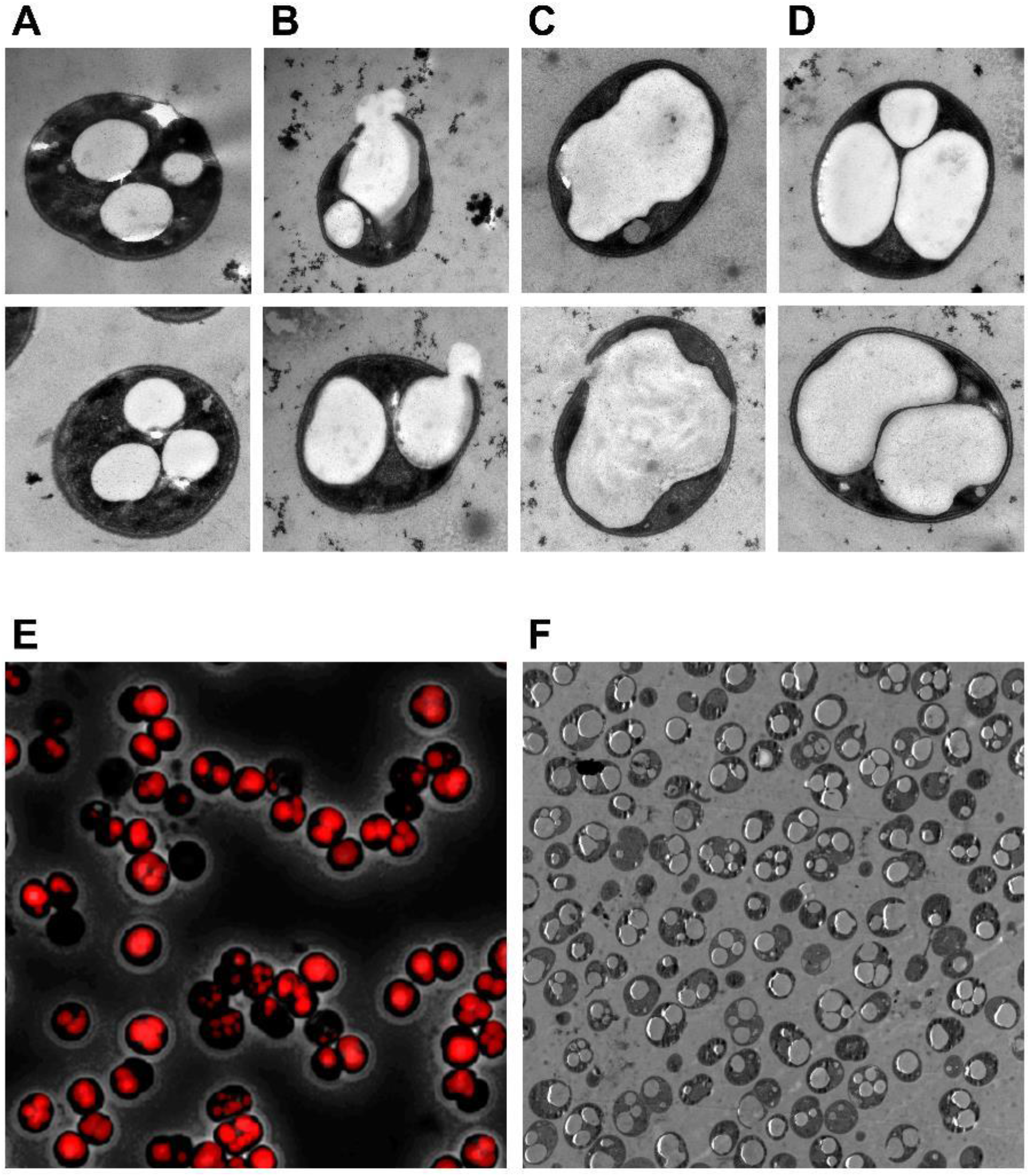
WT (A) and PPT1 cells (B-F) after 21 days of nitrogen-phosphorus-starvation with 10 mM acetate grown under alternating light/dark regime. (A) WT cells for comparison. (B) PTT1 cells showing a ruptured cell wall. (C) PPT1 cells with a single PHB granule. (D) PPT1 cells with multiple granules. (E) Fluorescence microscopic picture of PPT1 cells; PHB granules are visualized as red inclusions after staining with Nile red. (F) Overview of multiple PPT1 cells.

The samples for the images in Figure 5 were taken from the same cells, which were used for the experiment shown in Figure 4 B after 21 days (PPT1 cells, nitrogen-phosphorus starvation with 10 mM acetate). Electron-microscopic images show that the cells are fully packed with PHB granules (Figure 5 C, D). Although some heterogeneity among the cells is visible, most of the cells contained large quantities of PHB. The TEM pictures revealed that the interior of many cells was vastly filled up by PHB (Figure 5 C, D). Interestingly, most cells contained not multiple, but only one large PHB granule, indicating a potential fusion from smaller granules. In several cases, the observed cells were already ruptured, releasing PHB into their environment (Figure 5 B). In overview TEM pictures, it became apparent that most cells contained large quantities of PHB (Figure 5 F).

### Qualitative analysis of PHB

To further characterize the material properties of the produced PHB, PPT1 cells were cultivated for four weeks under nitrogen and phosphorus starvation. The cells were broken by sodium hypochlorite treatment and the purified PHB was analysed via gel permeation chromatography (GPC), 1H-NMR and 13C-NMR, to determine the molecular weight, the dispersity, the purity and the tacticity of the polymer, respectively. GPC analysis showed that PPT1 produces a high-molecular-weight polymer with relatively narrow dispersity (Figure 6). The number-average molecular weight was determined at Mn = 503 kg/mol (*Đ* = 1.74), which was more than twice as high than the control (Mn = 246 kg/mol, *Đ* = 2.33).

**Figure 6.**
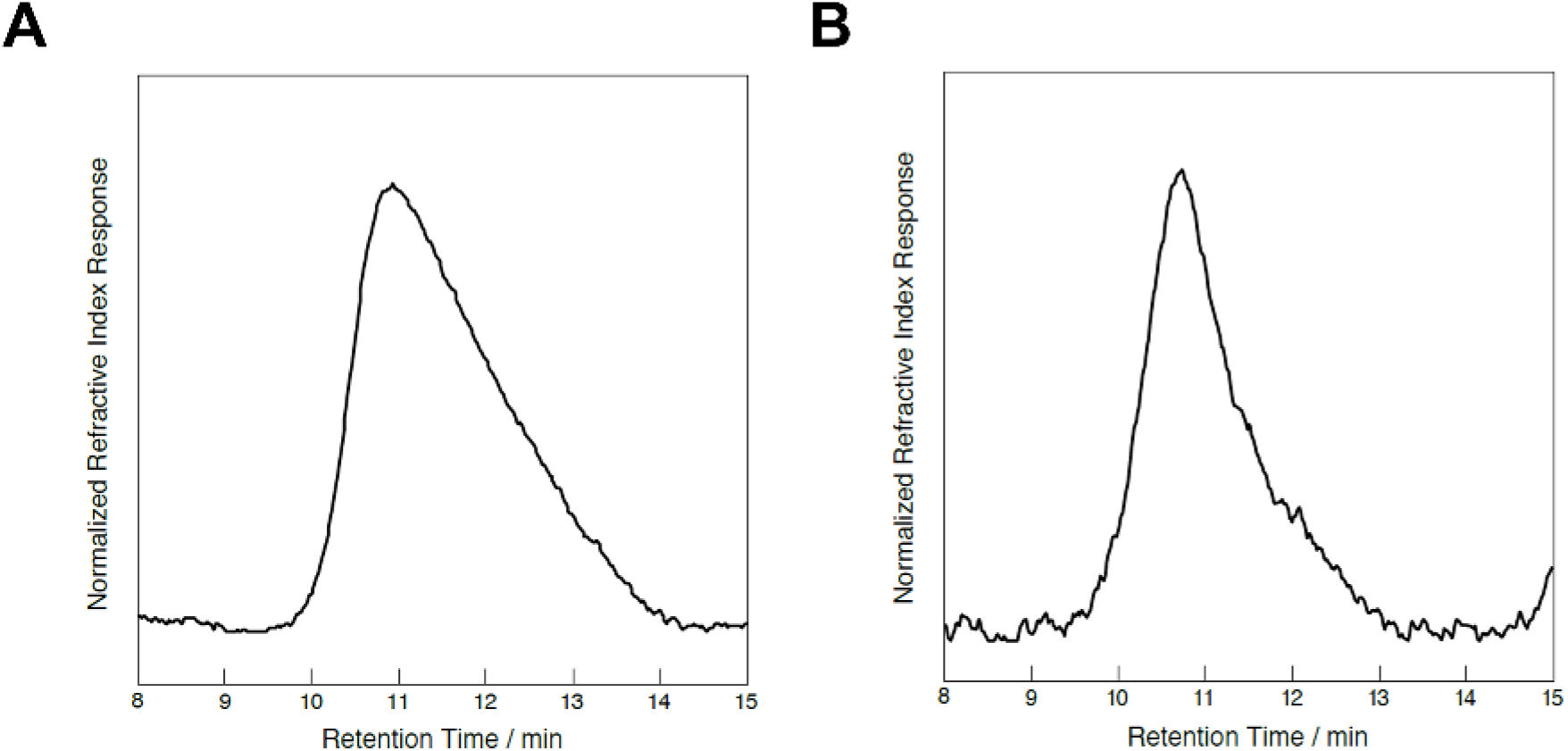
GPC analysis of PHB from an industrial standard (A) and PPT1 (B).

The chemical structure of the polymer was confirmed by 1H and 13C NMR spectroscopy to be completely pure PHB (Figure S6, S7). Furthermore, the observed singlet resonances in the 13C NMR spectrum indicated that the PHB derived from PPT1 is highly isotactic (Figure S8).

## 3. Discussion

### Δsll0944-REphaAB produces maximum amounts of PHB

As previous studies have shown, PHB is derived from the intracellular glycogen pools (Koch et al., 2019). Furthermore, this carbon flux is regulated by the protein Sll0944, which controls a central enzyme of the glycogen catabolism (phosphoglyceratemutase) (Orthwein et al., 2020). Deletion of Sll0944 results in strongly increased glycogen catabolism during prolonged nitrogen starvation. By the additional expression of the genes *phaA* and *phaB,* most of the carbon is redirected from the acetyl-CoA to the PHB pool. Since the reaction catalyzed by PhaB is converting one NADPH to NADP, the reaction yielding hydroxybutyryl-CoA is strongly favored during nitrogen starvation, where NADPH pools are increased (Hauf et al., 2013), driving PHB forward (Figure S1).

When grown in nutrient-replete balanced medium, the growth behavior of the PPT1 strain was comparable to the WT, in liquid medium as well as on solid agar plates (Figure 1, Figure S2). It is expected that both strains behave similar under these conditions, since there was hardly any PHB produced under vegetative grow (Figure S3) and since *sll0944* is mostly expressed during nitrogen starvation (Klotz et al., 2016b). The newly generated strain PPT1 shows a combinatory phenotype of both individual strains Δ*sll0944* and RE*phaAB*: while both individual strains have increased PHB production under nitrogen starvation by ~10 % / CDW, the strain PPT1 showed an additive phenotype of both effects and thereby reached values of up to 45 % / CDW (Figure 2 A). Similar PHB contents were reached regardless of the applied light regime, indicating that the production of PHB is not limited by the availability of light (Figure 2). The accumulation of PHB was further boosted by combined nitrogen-phosphorus starvation (Figure 3 C). This is in accordance with previous studies, where the combined nitrogen-phosphorus starvation caused the highest PHB production (Carpine et al., 2017). In contrast, the individual limitation of either sulfur or phosphorus resulted in only small intracellular PHB accumulations (Figure 3 A and B). It was shown before, that nitrogen limitation is most efficient for the induction of PHB synthesis in cyanobacteria (Kaewbai-Ngam et al., 2016). In a recently created strain though, it was shown that a random mutation in a phosphate specific membrane protein PstA caused a strong increase in PHB accumulation, hinting towards the importance of phosphorus for PHB production (Kamrava et al., 2018).

When 100 mM NaHCO_3_ were added to PPT1 cells cultivated in nitrogen-phosphorus depleted medium, a further increase of intracellular PHB levels was reached in the initial phase. This indicates that a limitation of carbon was impairing the PHB production in previous experiments. Since PHB is mostly formed from intracellular carbon (Dutt and Srivastava, 2018), carbon availability could be exhausted at such high PHB contents and thereby limit further accumulation of PHB. Notably, one of the three biological replicates exhibited a PHB content of 61 % / CDW after one week, indicating the potential to accelerate the pace of PHB formation by process engineering. The overall content was further increased by the addition of 10 mM acetate, hinting towards a limitation of the precursor acetyl-CoA. Since acetate can be converted to acetyl-CoA in a single enzymatic reaction, it is more efficiently metabolized to PHB compared to NaHCO_3_.

Interestingly, the highest PHB content was reached under light/dark regime, while its accumulation was strongly diminished under continuous light, even upon the addition of acetate (Figure 4 and Figure S4). This fits to previous observations, where cultivation under diurnal light/dark cycles was shown to increase the PHB production (Koch et al., 2020). Cells which were cultivated under conditions of gas-exchange limitation showed reduced PHB accumulation. This was also reported by other groups (Kamravamanesh et al., 2017) and might be explained by the lack of oxygen during the night, which is necessary for maintaining cell metabolism. Alternatively, excess of oxygen during the day could cause an increased oxygenase reaction which wastes energy and thereby slows down cell metabolism.

### Morphology of PHB granules

TEM pictures showed *Synechocystis* cells fully packed with PHB granules (Figure 5). Additionally, a certain number of cells displayed fractured cell envelops, leading to PHB granules leaking out of the cells. The rupture of cells could be due to intracellular mechanical pressure from the expanding PHB granules or it could be caused from mechanical stress during the preparation process. Whatever the cause of the ruptures was, it indicates an increased cell fragility due to the massive accumulation of PHB, since the effect was not detected in WT cells, which contained less PHB but were treated with the same procedure. This indicates that some PPT1 cells have reached an upper limit of how much PHB a cell can accumulate, above which cell viability is severely challenged. It was previously hypothesized that *Synechocystis* cells cannot accumulate larger quantities of PHB due to sterical hindrance of the thylakoid membranes. This study demonstrates that it is possible to manipulate *Synechocystis* in such a way that it accumulates vast amounts of PHB. Interestingly, most cells which contained large PHB-quantities possessed only very few granules, often just one single granule. This indicates that PHB granules merge together once they exceed a certain size.

### Qualitative analysis of PHB

Analysis of the extracted PHB derived from PPT1 revealed that it consists of PHB only. While other bacteria are able to produce PHAs with different side chains, such as 3-hydroxyvalaerate, the PhaEC enzyme, which is present in *Synechocystis*, is producing selectively PHB. For future experiments, a mutant strain harbouring a heterologous PHA polymerase could be created for the production of heteropolymers with improved material properties, such as poly(3-hydroxybutyrate-co-3-hydroxyvalerate) (PHBV). Other cyanobacteria, like *Nostoc microscopicum*, have already shown to possess PHA-polymerases which are able to produce PHBV (Tarawat et al., 2020). In previous analysis the average molecular weight of PHB from *Synechocystis* and *Synechocystis* sp. PCC 6714 was determined at Mn ~ 130 and 316 kg mol^−1^, respectively (Osanai et al., 2014, Lackner et al., 2019). Compared to this, the PHB derived from PPT1 is high-molecular, showing an average weight of 503 kg/mol. The PHB derived from PPT1 turned out to be highly isotactic, which is beneficial for good biodegradation properties.

### Conclusiosn and outlook

To further accelerate PHB production, overexpressing a strong PHB-polymerase could be beneficial. Although it was shown that higher levels of PhaEC can lower the PHB content (Khetkorn et al., 2016b), its activity could be rate limiting once such high values as in this present study are reached. The insertion of another short-chain-length PHA-polymerase could furthermore lead to the production of PHAs with improved material properties (PHBV). In order to improve the overall production yields, increased growth rates would be necessary, for example by the cultivation in high-density cultivators. In similar approaches, *Synechocystis* cultures reached OD_750_ of above 50 when higher light and CO_2_ concentrations were applied (Dienst et al., 2019, Lippi et al., 2018). Under those ideal conditions, up to 8 g of dry biomass l^−1^ d^−1^ were reached. If the time for chlorosis is assumed to be similar to the time required for cultivation and an intracellular PHB content of 60 % is reached, 2,4 g PHB l^−1^ d^−1^ could be produced under completely phototrophic conditions. Since the PHB production in the strain PPT1 is optimal under light/dark regime, the strain is also well suited for outdoor cultivation. Scaling up the cultivation to larger reactors would further reduce the production costs of PHB (Panuschka et al., 2019). Additionally, the ability of autotrophic cyanobacteria to sequester CO_2_ from the atmosphere could be beneficial for CO_2_ emission trading. Alternatively, a growth-coupled PHB production could be beneficial for certain kinds of cultivation

In summary, this study shows for the first time that cyanobacteria have the potential to accumulate large quantities of PHB. Furthermore, we demonstrate that also under cultivation with CO_2_ as the only carbon source, *Synechocystis* is able to produce quantities of PHB, which is of high relevance for the sustainable production of PHB as a bioplastic. This study helps to come closer to an industrial production of carbon neutral plastic alternatives.

## 5. Materials and Methods

### Cyanobacterial cultivation conditions

If not stated differently, *Synechocystis sp.* PCC 6803 cultures were grown in standard BG_11_ medium with the addition of 5 mM NaHCO_3_ (Rippka et al., 1979). The cultures were constantly shaken at 125 rpm, 28°C and at a illumination of ~50 μE. A 100 ml Erlenmeyer flask was used to grow 50 ml of bacterial culture. When cells were grown under alternating light/dark rhythm (12 hours each), the precultures were already adapted to these conditions by cultivating them under light/dark rhythm for two days. Whenever required, appropriate antibiotics were added to the mutant strains. When cultivation in depletion-medium was required, the following were used: for nitrogen starvation BG_0_ (BG_11_ without NaNO_3_); for sulfur starvation BG_11_ with MgCl instead of MgSO_4_; for phosphor starvation KCl instead of K_2_HPO_4_. Since *Synechocystis* has intracellular polyphosphate storage polymers, a preculture in phosphorus free medium was inoculated two days before the actual shift to phosphor free medium. For all starvations, exponentially grown cells (OD_750_ 0.4-0.8) were washed twice in the appropriate medium. For this, the cells were harvested at 4,000 g for 10 min, the supernatant discarded and the pellet resuspended in the appropriate medium. Afterwards the culture was adjusted to an OD_750_ of 0.4. For growth on solid surfaces, cells of an OD_750_ = 1 were dropped on BG_11_ plates containing 1.5 % agar. A serial dilution of the initial culture was prepared in order to count individual colony-forming-units. A list of used strains in this study is provided in Table 2.

**Table 2.**
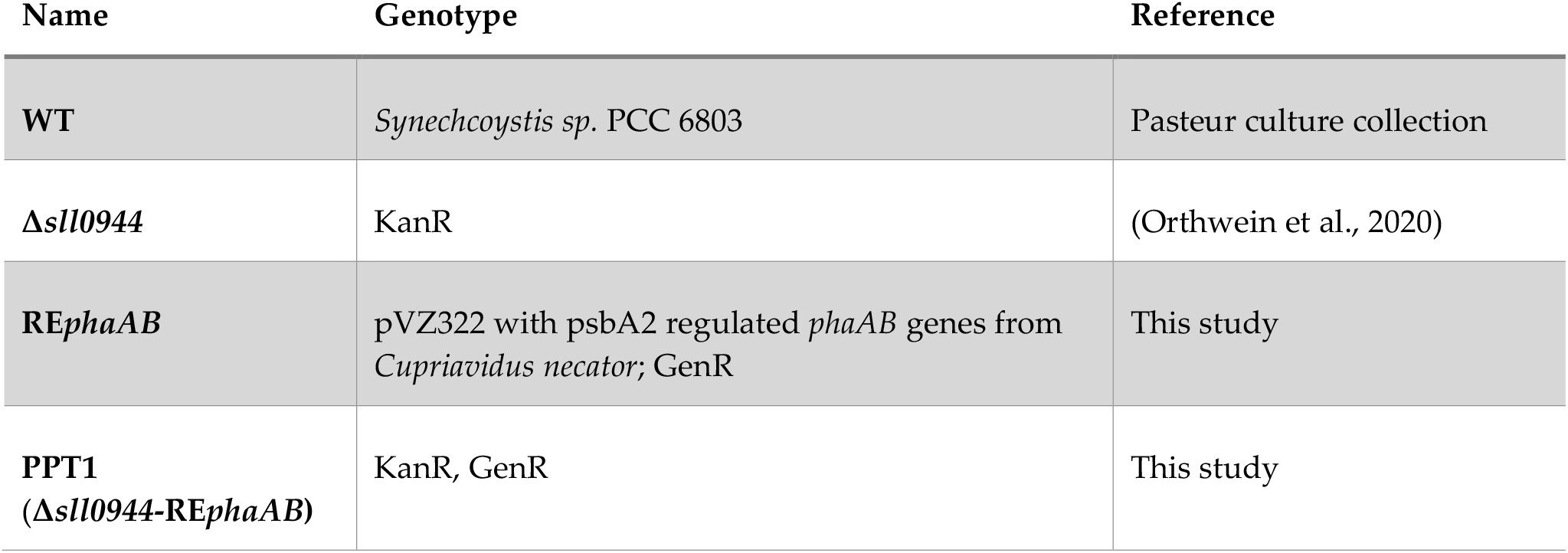
List of strains used in this study.

### Construction of RE*phaAB* and Δ*sll0944*-RE*phaAB* mutants

The promotor psbA2 and *phaAB* were amplified from genomic DNA of *Synechocystis* and C*upriavidus necator*, respectively. For this, the primer psbaA2fw2/psbA2rv2 or RephaABA2fw/RephaABA2rv were used (Table 3). A Q5 high-fidelity polymerase (NEB) was used to amplify the DNA fragments. The latter were subsequently assembled in pVZ322 vector (Gibson et al., 2009), which was beforehand opened at the XbaI site. The resulting vector was propagated in *E. coli* Top10 and isolated using a NEB miniprep kit. The plasmid was subsequently sequenced to verify sequence integrity. The correct plasmid was then transformed into *Synechocystis* using triparental mating (Wolk et al., 1984), resulting in the strain RE*phaAB*. The same RE*phaAB* plasmid was also transformed in the strain Δ*sll0944*, resulting in the strain PPT1 (Δ*sll0944*-RE*phaAB*).

**Table 3.**
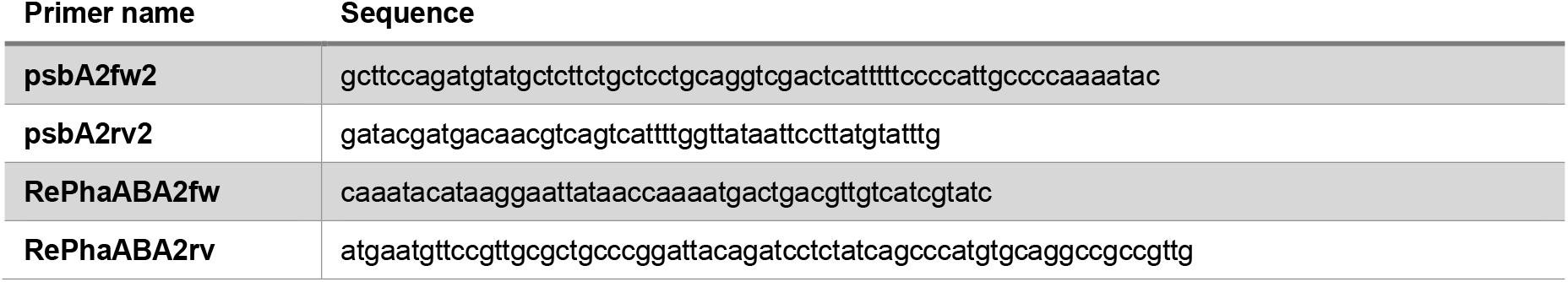
List of oligonucleotides used in this study.

### Gas exchange limitation

When gas exchange limitation was applied, 10 ml of culture were transferred to a 15 ml reaction tube. The tube was closed and additionally sealed with several layers of parafilm. During the incubation, the reaction tubes were constantly shaken.

### Microscopy and staining procedures

To analyze the intracellular PHB granules, 100 μl of *Synechocystis* culture were centrifuged (10,000 g, 2 min) and 80 μl of the supernatant discarded. Nile red (10 μl) was added and the pellet resuspended. From this mixture, 10 μl were dropped on an agarose-coated microscopy slide. For the detection, a Leica DM5500 B with an 100x /1.3 oil objective was used. An excitation filter BP 535/50 was used to detect Nile red stained granules.

### PHB quantification

To determine the intracellular PHB content, ~10 ml of cells were harvested by centrifugation (10 min at 4,000 g). The supernatant was discarded, and the remaining cell-pellet dried in a Speed-Vac for at least 2 h at 60°C. The weight of the dried pellet was measured to determine the CDW. Next, 1 ml of concentrated sulfuric acid (18 M H_2_SO_4_) was added and the sample was boiled for 1 h at 100°C. This process converts PHB to crotonic acid at a ratio of 1 to 0.893. The samples were diluted by transferring 100 μl to 900 μl of 14 mM H_2_SO_4_. Subsequently, the tubes were centrifuged for 10 min at 10,000 g. Next, 500 μl of the supernatant were transferred to a new tube and 500 μl of 14 mM H_2_SO_4_ were added. The samples were centrifuged again and 400 μl of the clear supernatant was transferred into a glass vile for HPLC analysis. For this, a 100 C 18 column (125 by 3 mm) was used with 20 mM phosphate buffer at pH 2.5 for the liquid phase. As a standard, a dilution series of commercially available crotonic acid was used. The final amount of crotonic acid was detected at 250 nm.

### Electron microscopy

For electron microscopic pictures, *Synechocystis* cells were fixed and post-fixed with glutaraldehyde and potassium permanganate, respectively. Subsequently, ultrathin sections were stained with lead citrate and uranyl acetate (Fiedler et al., 1998). The samples were then examined using a Philips Tecnai 10 electron microscope at 80 kHz.

### Purification of PHB

For the analysis of PHB, PPT1 cells were cultivated for four weeks in BG_11_ medium (without phosphorus and nitrogen) at light/dark regime. The cells were harvested by centrifugation for 10 min at 4,000 g. The cell pellet was resuspended in 15 ml freshly bought sodium hypochlorite solution (6 %) and shaken over night at room temperature. The next day, the cell debris were centrifuged and washed with water (10 times), until the chlorine smell disappeared. Subsequently, the pellet was washed once with 80 % ethanol and once with acetone.

### NMR and GPC

To characterize the chemical properties of PHB derived from PPT1, NMR spectra were recorded on a Bruker AVIII-400 spectrometer at ambient temperatures. As a control, an industrial standard PHB was used (BASF, Ludwigshafen, Germany). ^1^H and ^13^C NMR spectroscopic chemical shifts δ were referenced to internal residual solvent resonances and are reported as parts per million relative to tetramethylsilane. The tacticity of the polymer was analysed by ^13^C NMR spectroscopy according to literature (Bloembergen et al., 1989). As NMR solvent, CDCl_3_ was used (Sigma-Aldrich, Taufkirchen, Germany).

Measurements of polymer weight-average molecular weight (*M*_w_), number-average molecular weight (*M*_n_) and molecular weight distributions or dispersity indices (*Đ* = *M*_w_/ *M*_n_) were performed via gel permeation chromatography (GPC) relative to polystyrene standards on an PL-SEC 50 Plus instrument from Polymer Laboratories using a refractive index detector. The analysis was performed at ambient temperatures using chloroform as the eluent at a flow rate of 1.0 mL min^−1^.

## Author Contributions

Conceptualization, M.K. and K.F.; Methodology, M.K. and K.F.; Investigation, M.K.; Writing-Original Draft Preparation, M.K. and K.F.; Writing-Review & Editing, M.K and K.F.; Supervision, K.F.; Project Administration, M.K. and K.F.

## Acknowledgments

We thank Claudia Menzel for the preparation of the TEM pictures, Eva Nußbaum for the maintenance of cyanobacterial strains and technical assistance as well as Andreas Kulik for the operation of the HPLC.

