## Supplementary files for "Maximizing PHB content in *Synechocystis sp.* PCC 6803: development of a new photosynthetic overproduction strain"

**Supplementary Materials**

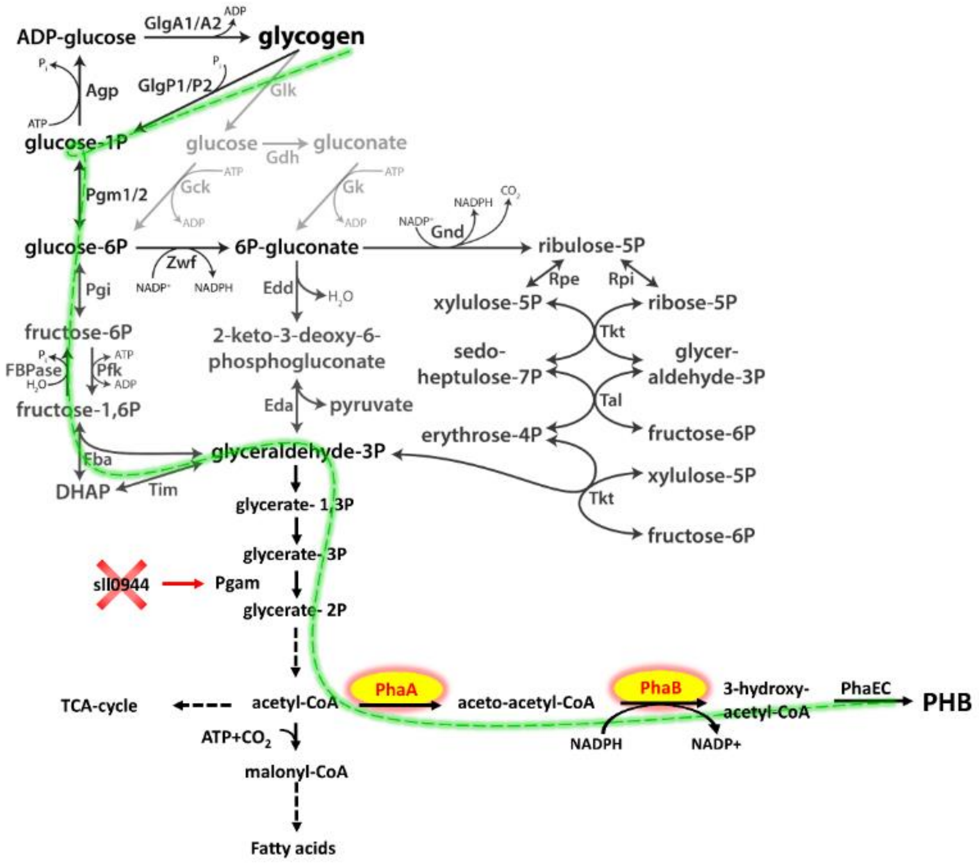


**Figure S1.** Overview about the central carbon-pathways in *Synechocystis*. It was previously shown, that the glycogen and PHB pool are interconnected via the EMP pathway (green) (Koch et al., 2019). Highlighted are the deleted protein Sll0944 and its target, the phosphoglyceratmutase, as well as the two overexpressed enzymes, PhaA and PhaB.

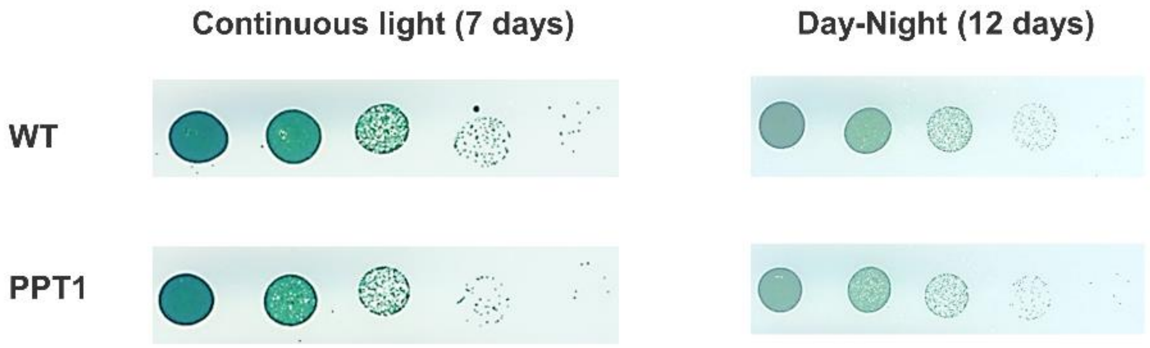


**Figure S2.** Drop plate assay of the WT and PPT1. Vegetative cells at an $OD_{750}$ of 1 were diluted 10-fold for five times ($10^0$ to $10^4$, respectively). The dilutions were then dropped on a $BG_{11}$ agar plate and grown at continuous light or light/dark rhythm for 7 or 12 days, respectively. The plate shown in the figure is representative for 3 individually grown biological replicates.

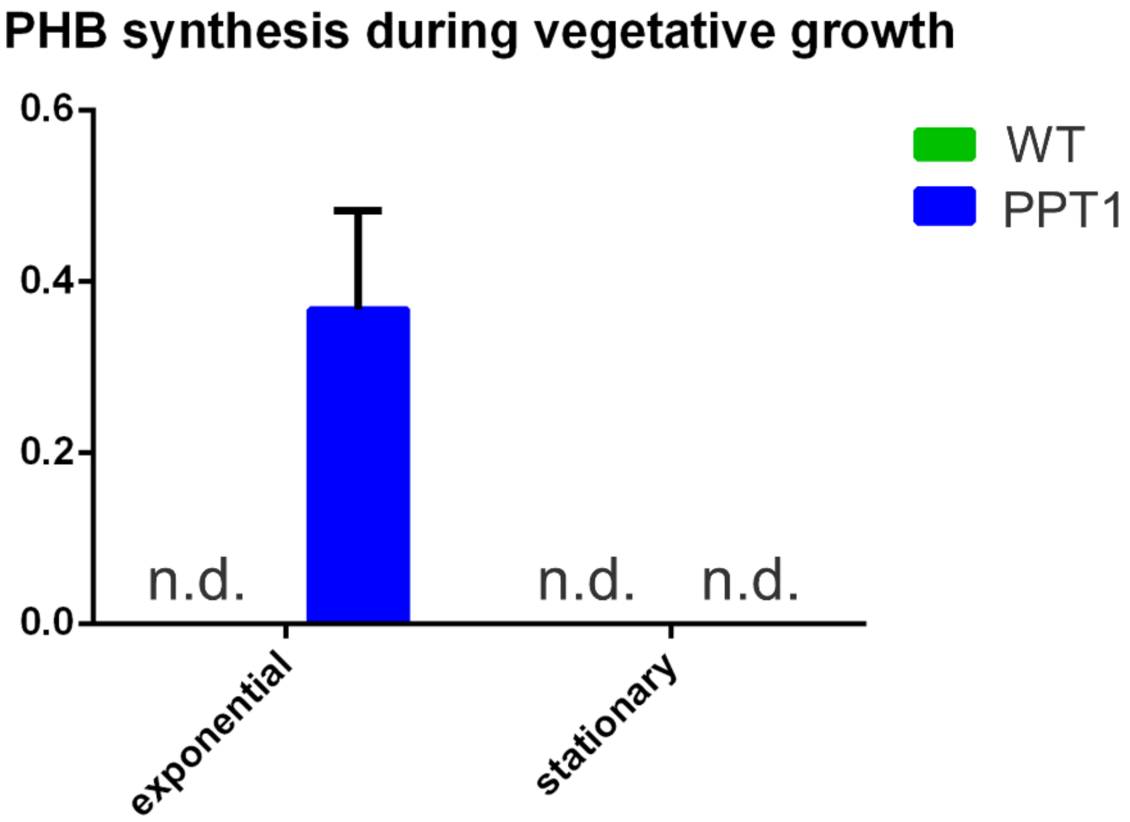


**Figure S3.** PHB accumulation under vegetative growth. WT and PPT1 cells were sampled during exponential or stationary phase (OD ~1 and ~3, respectively) at continuous lighting. n.d. = not detectable. Each point represents a mean of three independent biological replicates.

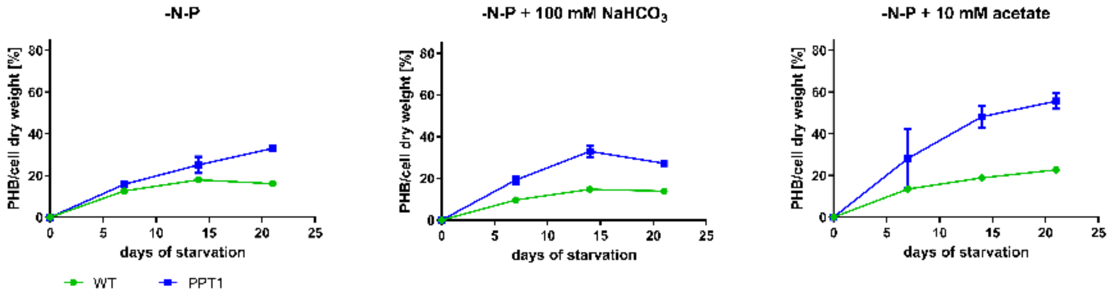


**Figure S4.** PHB production of WT (green) and PPT1 (blue) cells grown under continuous lighting. Cells shifted to nitrogen/phosphor free medium (A) and with additional 100 mM $NaHCO_3$ (B) or 10 mM acetate (C). Each point represents a mean of three independent biological replicates.

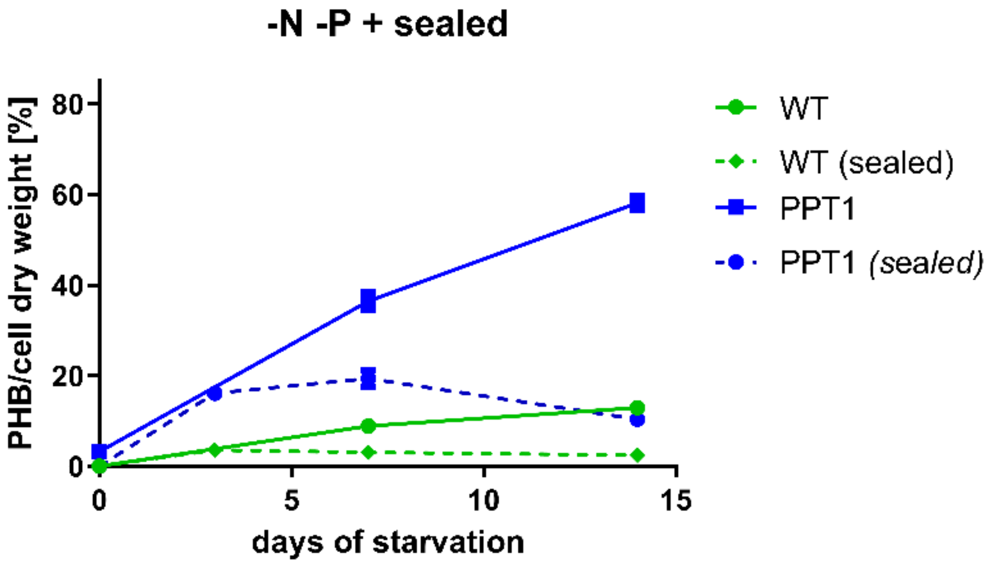


**Figure S5.** PHB content of WT (green) and PPT1 (blue) cells grown in nitrogen/phosphor free medium under light/dark regime. Dashed lines indicate growth in sealed vessels. Each point represents a mean of three independent biological replicates.

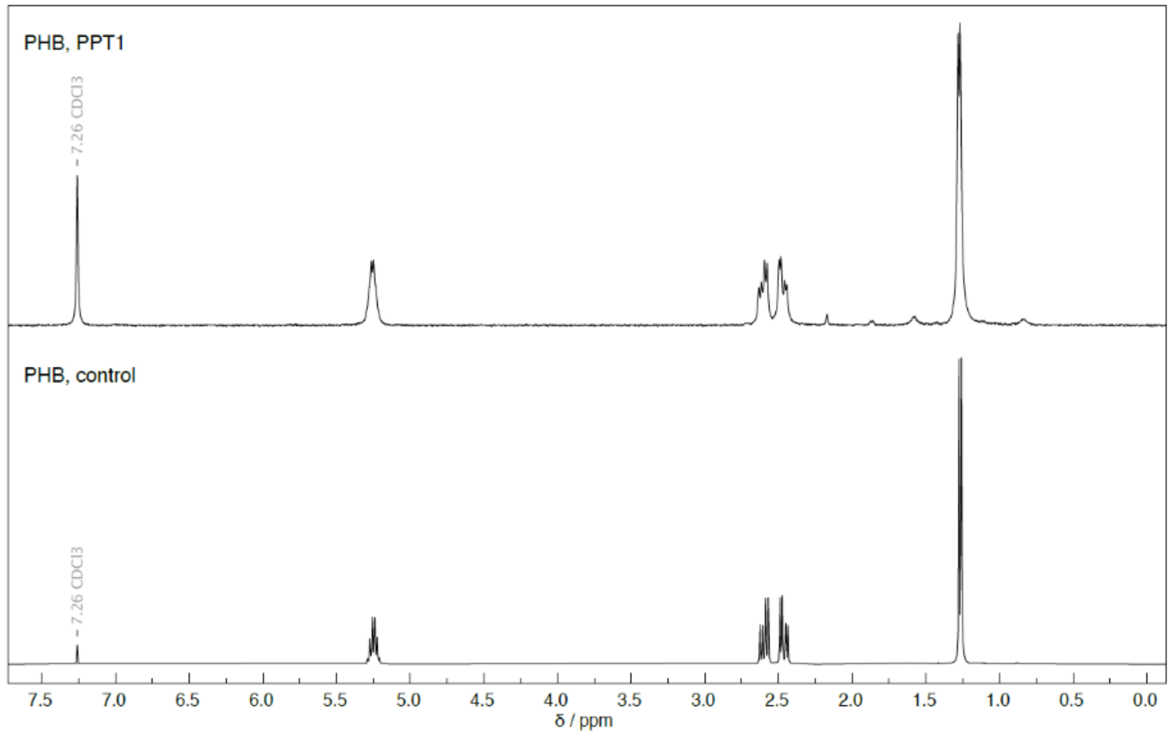


**Figure S6.** $^1H$ NMR ($CDCl_3$, 400 MHz) spectrum of PHB derived from PPT1 compared to an industrial standard sample.

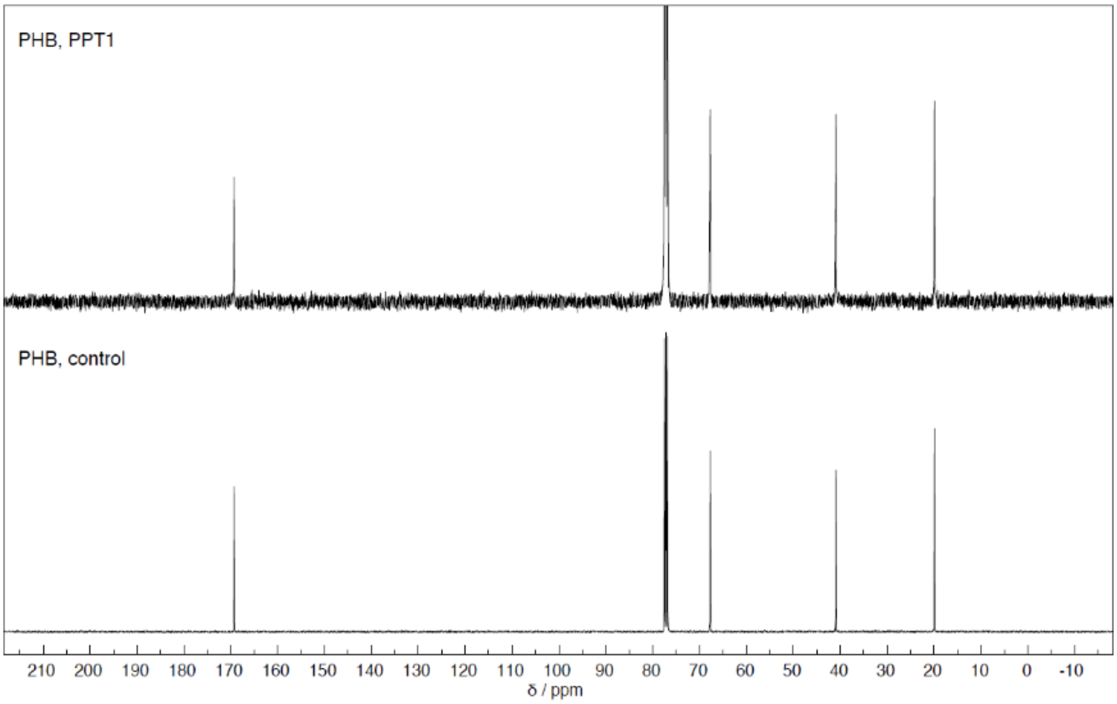


**Figure S7.** [13]C NMR spectrum ($CDCl_3$, 101 MHz) of PHB derived from PPT1 compared to an industrial standard sample.

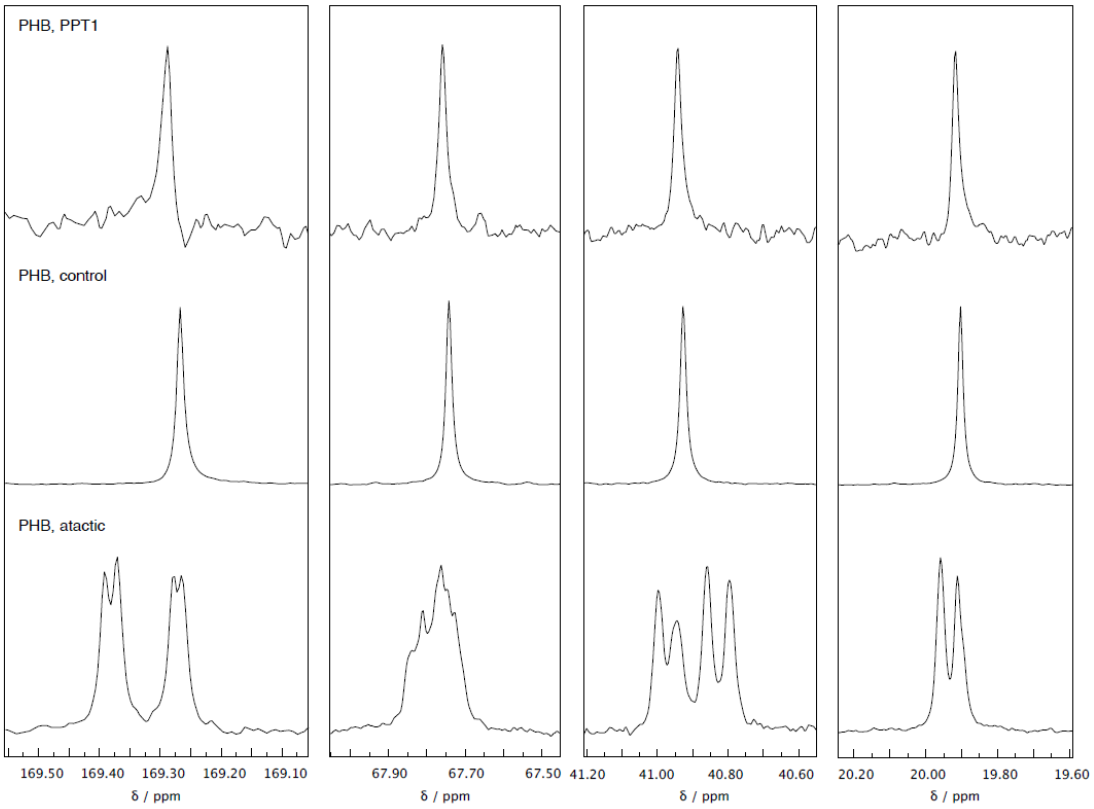


**Figure S8.** [13]C NMR spectrum to analyse the tacticity of PHB derived from PPT1. For comparison, industrial standard PHB (isotactic) and atactic PHB (produced from ß-butyrolactone via ring-opening polymerization) are shown.
